# Lack of evidence for participation of TMEM150c/TENTONIN3 in sensory mechanotransduction

**DOI:** 10.1101/2022.02.13.478866

**Authors:** Julia Ojeda-Alonso, Valérie Bégay, Jonathan Alexis Garcia-Contreras, Andrea Fernanda Campos-Pérez, Bettina Purfürst, Gary R. Lewin

## Abstract

The membrane protein TMEM150c has been proposed to form a mechanosensitive ion channel that is required for normal proprioceptor function. Here we examined whether expression of TMEM150c in neuroblastoma cells lacking Piezo1 is associated with the appearance of mechanosensitive currents. Using three different modes of mechanical stimuli, indentation, membrane stretch and substrate deflection we could not evoke mechanosensitive currents in cells expressing TMEM150c. We next asked if TMEM150c is necessary for the normal mechanosensitivity of cutaneous sensory neurons. We used an available mouse model in which the *Tmem150c* locus was disrupted through the insertion of a LacZ cassette with a splice acceptor that should lead to transcript truncation. Analysis of these mice indicated that ablation of the *Tmem150c* gene was not complete in sensory neurons of the dorsal root ganglia (DRG). Using a Crispr/cas9 strategy we made a second mouse model in which a large part of the *Tmem150c* gene was deleted and established that these *Tmem150c*^-/-^ mice completely lack TMEM150c protein in the DRGs. We used an *ex vivo* skin nerve preparation to characterize the mechanosenstivity of mechanoreceptors and nociceptors in the glabrous skin of the *Tmem150c*^-/-^ mice. We found no quantitative alterations in the physiological properties of any type of cutaneous sensory fiber in *Tmem150c*^-/-^ mice. Since it has been claimed that TMEM150c is required for normal proprioceptor function we made a quantitative analysis of locomotion in *Tmem150c*^-/-^ mice. Here again we found no indication that there was altered gait in *Tmem150c*^-/-^ mice compared to wild type controls. In summary, we conclude that existing mouse models that have been used to investigate TMEM150c function in vivo are problematic. Furthermore, we could find no evidence that TMEM150c forms a mechanosensitive channel or that it is necessary for the normal mechanosensitivity of cutaneous sensory neurons.

## Introduction

The search for genes coding for proteins that are necessary for touch driven behavior was pioneered using *C. elegans* by Martin Chalfie and colleagues (Arnadóttir and Chalfie, 2010). These studies demonstrated how touch neurons transduce mechanical stimuli with a core mechanosensitive ion channel that works in the context of its interaction with several regulatory proteins that are themselves also necessary for mechanotransduction. Thus, the central MEC4/MEC10 ion channel conducts the ionic currents initiated by force (O’Hagan et al., 2005; Driscoll and Chalfie, 1991), but requires accessory proteins like MEC2 and MEC6 in order to function (O’Hagan et al., 2005; Goodman et al., 2002; Huang et al., 1995; Chelur et al., 2002; Brown et al., 2008). In recent years there has been considerable progress in identifying the molecules involved in force transduction by mammalian touch receptors. Thus, the MEC-2 like protein STOML3 was shown early on to be necessary for mammalian mechanoreceptors to transduce mechanical force into an electrical signal, a process we have termed sensory mechanotransduction (Wetzel et al., 2017, 2007). It is less clear whether mammalian orthologues of MEC4/MEC10 such as members of the Acid Sensing Ion Channel family (ASICs) participate directly in the transduction of mechanical force in sensory neurons (Omerbašić et al., 2014). The discovery of PIEZO1 and PIEZO2, very large membrane proteins that can form homo-trimeric ion channels opened by mechanical force (Coste et al., 2010), raised the possibility that such proteins may participate in sensory mechanotransduction. Indeed, conditional deletion of the *Piezo2* gene in mouse sensory neurons leads to the silencing of around half of all cutaneous mechanoreceptors (Ranade et al., 2014; Murthy et al., 2018), a phenotype that is almost identical to that observed in *Stoml3^-/-^* mice (Wetzel et al., 2017, 2007). Indeed, we have shown that STOML3 is a powerful regulator of PIEZO channel sensitivity to mechanical force (Poole et al., 2014a). Analysis of mice in which the *Piezo2* or *Stoml3* genes were ablated made it clear that the mechanosensitivity of many mechanoreceptors and almost all nociceptors relies on as yet unknown mechanosensitive ion channels. This fact has led many groups to search for further ion channel candidates that could confer mechanosensitivity to sensory neurons (Patkunarajah et al., 2020; Beaulieu-Laroche et al., 2020; Hong et al., 2016; Parpaite et al., 2021).

One early candidate sensory transduction channel was proposed to be the membrane protein TMEM150c, which has 6 predicted transmembrane domains (Hong et al., 2016). Subsequent studies suggested that the mechanosensitive currents identified after expression of TMEM150c in Hek293 cells were due to the confounding presence of endogenous PIEZO1 channels (Dubin et al., 2017; Anderson et al., 2018). However, one study provided evidence that TMEM150c might modulate the kinetics of PIEZO2 channel activation (Anderson et al., 2018). The acquisition of mechanosensitivity by sensory neurons in the form of mechanosensitive currents is a developmentally regulated process occurring as soon as sensory axons reach their peripheral targets (Lechner et al., 2009). We had used this phenomenon to screen for the induction of genes encoding transmembrane proteins in sensory neurons coincident with the appearance of mechanosensitivity. One gene that was highly induced coincident with target innervation was *Tmem150c* (Herget and Lewin, unpublished data), a finding that also prompted us to investigate the role of this gene in cutaneous mechanosensation. We generated mutant mice with *Tmem150c* gene ablation and examined these mice for sensory deficits. If TMEM150c, is itself an ion channel or, like STOML3, a necessary modulatory subunit involved in sensory mechanotransduction, we would expect to find deficits in the physiology of cutaneous sensory neurons. Instead, after a detailed physiological evaluation of two mutant mouse models with an ablated *Tmem150c* locus we conclude that there is no good evidence that this gene is involved in sensory mechanotransduction.

## Materials and Methods

### Generation of the *Tmem150c* mutant mice and genotyping

Design and generation of guide RNAs used for targeting CrispR/cas9 region of *Tmem150c/Ttn3* locus in pronucleus of C57Bl/6N, generation of founder mice and F1 heterozygous mice were performed by Ingenious Targeting Laboratory (Ingenious Targeting Laboratory, Ronkonkoma, NY). Briefly, the following primers were used to amplify the wild-type allele (Tmem150c WT Rev1: 5’-TACCTGATGTATGGAGCATGCTTC-3’ and Tmem150c WT Fw1: 5’-TACTTTATAGCCGTGGAAGATGAC-3’) and the mutant allele (MEMT 1 Fw: 5’-CTCAATAACAGCCACAAGGAAAG-3’ and MEMT 1 Rev: 5’-ACTGGCAGGGTTGTGTAAG-3’). F1 founders and F2 *Tmem150c*^+/-^ mice were crossed with C57Bl/6N mice. F3 and F4 heterozygous mice were mated to generate homozygous progenies used in the experiments. Progenies of heterozygous matings were born with Mendelian frequency that did not differ significantly from that expected (genotyped newborns n= 116; +/+ 25.9%: +/− 56%: −/− 18.1)

### *Tmem150c KOMP* line and genotyping

*Tmem150c^+/LacZ^* mice were generated from an ES cell clone created by international knockout mouse project (www.komp.org)(Tate and Skarnes, 2011). The genotyping of the progenies were performed according to KOMP protocol. Briefly, the following primers were used to amplify a wild-type band (forward primer 5’-CAC ATT GGC AAT CAG CAT TAC AC-3’ and reverse primer 5’-GTG CTG GGA TCC TCC ATT ACC-3’), a part of the neomycin cassette (CDS-neoF 5’-GGG ATC TCA TGC TGG AGT TCT TCG-3’ and the above reverse), and a part of the LacZ cassette (LacZ-F 5’-ATC ACG ACG CGC TGT ATC-3’ and LacZ-R 5’-ACA TCG GGC AAA TAA TAT CG-3’).

### RNA extraction and Real-Time PCR

Total RNA was isolated from tissues (epididymis, lumbal DRGs and liver) using Relia prep™ RNA Tissue Miniprep System (Promega) according to the manufacturer’s protocol. Total RNA was reverse transcribed at 42°C by using GoScript Transcription System (Promega) according to the manufacturer’s protocol. Real-time-PCR (RT-PCR) were run on a BioRad cycler. The following primers were used to amplify *Tmem150c* Ex4-Ex6 fragment (forward primer: 5’-GAAGCATGCTCCATACA-3’and reverse primer 5’-CCAAGTAAGGTCATTCC-3’), Ex7-Ex8 fragment (forward primer: 5’-AGCTCACAAACGATGAA-3’ and reverse primer: 5’-CAGGATGAAATACAGGA-3’) and *Hprt1* fragment (forward primer: 5’TCCTCCTCAGACCGCTTTT-3’ and reverse primer 5’-CCTGGTTCATCATCGCTAATC-3’).

### X-gal staining

Tissues were dissected and quickly fixed in 2% PFA, 0.2% glutaraldehyde in 0.1 M sodium Phosphate buffer for 3 hr and postfixed with X-gal fixative (0.2% glutaraldehyde, 100mM MgCl_2_, 50 mM EDTA in PBS) for 20 min. Postfixed lumbar DRGs and epididymis were washed in PBS and cryoprotected in 30% sucrose overnight at 4°C. Tissues were embedded in O.C.T Tissue Tek (Sakura Finetek, Netherland) on dry ice and stored at −80°C. Sections were cut at 12 □m using a Cryostat (Thermofisher) and X-galactosidase (X-gal) staining was performed as previously described (Lobe et al., 1999). Briefly, sections were incubated in X-gal reaction buffer (5 mM potassium ferrocyanide, 5 mM potassium ferricyanide, 2 mM MgCl2, 0.02% Nonidet P-40, 0.01% sodium deoxycholate and 0.5 mg/ml of X-gal in 100 mM sodium Phosphate, pH 7.3) for 1hr (for epididymis) to overnight (DRGs) at 37°C. Sections were washed in PBS and counter-stained with nuclear fast-red (N8002 Sigma). Sections were observed and captured on a Leica DM 5000 B microscope (Leica, Germany).

### Immunoblotting

Tissues from *Tmem150c*^-/-^ mutants and wild type mice were lysed with 8 M urea buffer and protein concentrations were determined using the Bradford reagent (Sigma). Proteins were separated by SDS-PAGE, followed by western blotting using rabbit antibody against the C-terminal part of TMEM150c (ABN2266, Millipore), mouse β-actin (A1978, Sigma), and horseradish peroxidase-conjugated secondary antibodies (111-035-003, Jackson Immuno-research), and chemiluminescence detection (Thermo Fischer).

### Electron microscopy

The general procedure followed for the quantification of peripheral nerves was described by (Smith et al., 2012). Tibial nerves from 30 week-old WT and *Tmem150c^-/-^* mice were analyzed. Nerves were freshly isolated and fixed in 4% paraformaldehyde/2.5% glutaraldehyde in 0.1 M phosphate buffer for at least 24 hr at 4°C. After treatment with 1% OsO4 for 2 hr, each nerve was dehydrated in a graded ethanol series and propylene oxide and embedded in polyresin. Nerves were cut using a microtome in semi-thin (1 μm) and ultrathin sections (70 nm). Semi-thin sections were stained with toluidine blue and ultrathin sections were contrasted with uranyl acetate and lead acetate. Semi-thin sections were examined in a light microscope to determine the total number of myelinated axons within the nerve. Ultrathin sections were examined with a Zeiss 910 electron microscope and images were taken at an original magnification of 2500x. Fifteen pictures (22.5 x 14.9 μm) per ultrathin section from each nerve from three adult WT and *Tmem150c*^-/-^ mice were analyzed. Using iTEM software, the C:A-fibers ratio, the myelin thickness and g-ratio of the myelin sheath of A-fibers, were determined. The g-ratio is defined as the ratio of the inner axonal diameter to the total outer diameter of the myelin sheath.

Based on the C:A-fibers ratio determined in ultrathin sections, the total number of C-fibers within the nerve was extrapolated to the number of A-fibers of each semi-thin section. To further compare between different animals, the total number of myelinated and unmyelinated axons was extrapolated to the smallest nerve area.

#### Cell culture and transfection

DNA plasmids that contain mouse *Tmem150c, mousePiezo1* and *mousePiezo2* sequences were obtained using large-scale bacterial culture (200 ml maxiprep). Single colonies from transformed bacteria were picked into LB medium containing (ampicillin or kanamycin), grown overnight at 37 °C with shaking. The maxi-preps were carried out according to the manufacture’s protocol. Plasmid DNA was diluted in water and the DNA concentration was measured using the NanoDrop2000 spectrophotometer (Thermo Fisher Scientific, USA).

N2a cells (N2a) from mouse were used. N2a cells present endogenous mechanically gated currents due to the expression of Piezo1. For that reason, in most of the experiments N2a cells where Piezo1 was deleted (N2a-P1KO) via CRISPR/Cas technology were used (Moroni et al., 2018). Cells were plated onto PLL glass coverslips or PLL covered pillar arrays in medium without serum and let them attach for at least 4 hours. For the transfection, 1 μg of DNA was mixed with 3 μl of FuGeneHD and 300 μl of OptiMem. After 15 min of incubation at room temperature, the mixture was added to each well. Cells were incubated at 37 °C in 5% CO_2_ overnight. Efficiently transfected cells could be detected by a fluorescent marker signal at least 24 hr after transfection.

### Whole-cell electrophysiology

Whole-cell recordings in voltage clamp were performed from the N2A cell line PIEZO1 knockout. Patch pipettes were pulled from borosilicate glass with a tip resistance of 3–5 MΩ were filled with intracellular solution (in mM): 100 KCl, 10 NaCl,1 MgCl2, 10 HEPES, and pH adjusted to 7.3 with KOH. The extracellular solution contained (in mM): 140 NaCl, 4 KCl, 1 MgCl2, 2CaCl2, 4 glucose and 10 HEPES. Cells were clamped to a holding potential of −60 mV. Recordings were made using an EPC-10 amplifier (HEKA, Germany) with Patchmaster and Fitmaster software (HEKA). Pippette and membrane capacitance were compensated using the auto function of Patchmaster and series resistance wascompensated by 70% to minimize voltage errors.

### Indentation

Mechanically activated currents were recorded as described earlier (Hu and Lewin, 2006; Wetzel et al., 2007). The mechanical probe consisted of fire-polished glass pipette (tip size 2-3μm) was manipulated by a piezo-driven micromanipulator (Nanomotor MM3A; Kleindiek Nanotechnik). Calibration of a single-step size was achieved moving a large number of steps and measuring the distance, repeated three times. Motor’s velocity of movement was set at 3.5 μm/ms. The voltage signal sent to the nanomotor by the control unit was simultaneously monitored by a second channel at the EPC10 amplifier, allowing to measure the delay between the nanomotor movement and mechanically activated current, or latency. The probe was positioned at 60° near to the cell body and movements of the mechanical probe were executed in the in/out axis of the device (axis C) for 300-500ms with 500ms pause in between steps. A voltage divider box (MM3-LS trigger, Kleindiek Nanotechnik) was connected that provided analog voltage output signal corresponding to the piezo signals. Depending on the movement of piezo micromanipulator, the voltage signal was registered as small pulse or just a change in a DC voltage. Only the somas were stimulated, the evoked currents were recorded with a sampling frequency of 200kHz.

### High Speed Pressure Clamp

High Speed Pressure Clamp (HSPC) recordings were performed in excised outside-out patches pulled from N2a cells at room temperature. Recording pipettes were prepared using a DMZ puller and subsequently polished to a final resistance of 6–8MΩ for outside-out patches. Currents were elicited by negative pressure stimuli, with an Ala Instrument device, applied through the recording pipette holding at −60mV. A protocol of pressure steps from 10mmHg to 150mmHg in 20 mmHg steps during 600ms was used. Uncompensated series resistance values were less than 2MΩ. The recorded currents responsive to the pressure curve from each cell were fitted to a Boltzmann equation using the FitMaster program (HEKA, Elektronik GmbH, Germany).

### Pillar arrays

Pillar arrays were prepared as previously described (Patkunarajah et al., 2020; Poole et al., 2014a; Servin-Vences et al., 2017). Briefly, silanized negative masters were used as templates. Negative masters were covered with polydimethylsilozane (PDMS, syligard 184 silicone elastomer kit, Dow Corning Corporation) mixed with a curing agent at 10:1 ratio. After 30 min, glass coverslips (thickness 2) were placed on the top of the negative masters containing PDMS and the coated master placed at 110°C for 1 hr. After curing, the pillar array was peeled away from the master. The resulting radius- and length-size of individual pilus within the array was 1.79 μm and 5.8 μm, respectively. While the elasticity and the spring constant of each pilus was 2.1 MPa and 251 pN-nm, respectively, as previously reported (Patkunarajah et al., 2020; Poole et al., 2014; Servin-Vences et al., 2017). Before use, pillar arrays were activated by plasma cleaning and coated with PLL (Deiner Electronic GmbH, Germany) and cells were allowed to attach.

To generate quantitative data an individual pilus subjacent to a cell was deflected using a heat-polished borosilicate glass pipette (approx. 2 mm in diameter) driven by a MM3A micromanipulator (Kleindiek Nanotechnik, Germany) as described in (Poole et al., 2014; Servin-Vences et al., 2017). Pillar deflection stimuli were applied in the range of 0-1000 nm and the electrical response of the cells was monitored using whole-cell patch-clamp. A bright-field image (Zeiss Axio Observer A1 inverted microscope) was taken before, during and after pillar deflection stimuli using a 40x objective and a CoolSnapEZ camera (Photometrics, Tucson, AZ). The pillar movement was calculated comparing the light intensity of the center of each pilus before and after the stimuli with a 2D-Gaussian fit (Igor Software, WaveMetrics, USA).

### Extracellular recording from tibial nerve

Electrophysiological recordings from cutaneous sensory fibers of the tibial nerve were made using an *ex vivo* skin nerve preparation following the method described previously (Walcher et al., 2018; Wetzel et al., 2007). Briefly, the animal was sacrificed by cervical dislocation and the hair of the limb was shaved off. The glabrous skin from the hind paw was removed along with the tibial nerve dissected up to the hip and cut. The glabrous skin along with the tibial nerve still attached to the hindpaw was transferred to a bath chamber which was constantly perfused with warm (32°C) oxygen-saturated interstitial fluid. The remaining bones, muscle and ligament tissue were gently removed as much as possible, allowing the glabrous skin and tibial nerve preparation to last at least 6 hours of recording in healthy a stable condition in an outside-out configuration. The tibial nerve was passed through a narrow channel to an adjacent recording chamber filled with mineral oil.

Single-unit recordings were made as previously described (Walcher et al., 2018; Wetzel et al., 2007). Fine forceps were used to remove the perineurium and fine nerve bundles were teased and placed on a platinum wire recording electrode. Mechanical sensitive units were first located using blunt stimuli applied with a glass rod. The spike pattern and the sensitivity to stimulus velocity were used to classify the unit as previously described (Walcher et al., 2018; Wetzel et al., 2007). Raw data were recorded using an analog output from a Neurolog amplifier, filtered and digitized using a Powerlab 4/30 system and Labchart 8 software with the spike-histogram extension (ADInstruments Ltd., Dunedin, New Zealand). All mechanical responses analyzed were corrected for the latency delay between the electrical stimulus and the arrival of the action potential at the electrode. The conduction velocity (CV) was measuring the formula CV = distance/time delay, in which CVs > 10ms^-1^ were classified as RAMs or SAMs (Aβ, <10ms^-1^ as Aδ and <1ms^-1^ as C-fibers.

#### Mechanical Stimulation

Mechanical stimulation of the receptor field of the recorded fibers was performed using a piezo actuator (Physik Instrumente, Germany, P-602.508) connected to a force measurement device (Kleindiek Nanotechnik, Reutlingen, Germany, PL-FMS-LS). Different mechanical stimulation protocols were used to identify and characterize the sensory afferents. Mechanoreceptors were tested with a vibrating stimulus with increasing amplitude and 20 Hz frequency. The force needed to evoke the first action potential was measured. Additionally, a ramp and hold step was used with Constant force (100mN) and repeated with varying probe movement velocity (0.075, 0.15, 0.45, 1.5 and 15mm s^-1^). Only the firing activity evoked during the dynamic phase were analyzed. SAM mechanoreceptors and nociceptors were tested with a mechanical stimulus with a constant ramp (1.5-2 mN ms^-1^) and increasing force amplitude, spikes evoked during the static phase were analyzed.

### Gait Analysis

Gait analysis was carried out using the “Mouse walk” method described by Mendes and colleagues (Mendes et al., 2015). An acrylic walkway 80cm in length was produced with white-light LED-strips attached to its sides. Another similar walkway was used as a background light (sky) with white plastic attached to the top to provide a lighter and even background. These two walkways were embedded in a black custom-built rack with a mirror angled 45° below the walkway. On the left side of the walkway a small custom-built plexiglas “house” was introduced to provide shelter for the animals. A monochromatic high-speed camera (FLIR, RoHS 2.3 MP Mono Grasshopper) was used to videotape the animal’s walk (the mirror image / reflection was recorded). Experiments were performed with no additional light source present in the room. Animals were initially habituated to the walkway for 15 minutes – they were put in the setup on the right side, opposite of the shelter and allowed to explore freely. Usually they spent most of the time in the shelter after the initial exploration of the walkway. At the end of the habituation time the shelter containing the animal was removed, the animal placed on the right side again and the shelter placed on the left side – to make the animal cross the walkway a few times. Each animal was recorded multiple times to obtain recordings with constant velocity and no stops during the crossing – 4 to 5 of these videos were then cropped, converted to single image stacks (every frame was converted into 1 image, the framerate of the camera was 150 frames/s) and analyzed using a MATLAB script provided by Cesar Mendes. The white-LED light transilluminates the walkway creating an effect called fTIR (frustrated total internal reflection) - in brief: the light traveling through the walkway is reflected differently when there is contact of the animals’ paw with the walkway. This is visualized through lighting of the contact area – the intensity of the reflected footprint is increasing with pressure / weight. This was used for the annotation of the videos, the paws as well as the head, nose, center of the body and other markers were used to evaluate approximately 26 parameters of the animals’ gait.

## Results

Initial reports suggested that heterologous expression of *tmem150c* is associated with the appearance of mechanosensitive currents in Hek293 cells to indentation (Hong et al., 2016). Mechanosensitive currents can be measured using a variety of different methods to apply mechanical force. The most commonly used method is to poke cells that express the proteins of interest, but currents can also be evoked by applying fast pressure pulses to patches of membranes in cell attached mode (Taberner et al., 2019) or in excised outside out patches (Moroni et al., 2018). In addition, some ion channels such as TRPV4 and PIEZO2 are not efficiently gated by membrane stretch, but are activated by displacement of the extracellular matrix underneath cells expressing the channel protein (Poole et al., 2014b; Servin-Vences et al., 2017). Here we expressed *tmem150c* in N2A cells in which the PIEZO1 gene had been ablated (N2A^*Piezo1*-/-^ cells) (Moroni et al., 2018) and evaluated whether mechanosensitive currents could be evoked with all three of the methods described above. In agreement with studies using Hek293 cells lacking PIEZO1, we found no poking-induced currents in N2A^*Piezo1*-/-^ cells transfected with a *tmem150c* cDNA similarly to the vector control (Figure 1A,B). In contrast, in all cells recorded expressing wild type *piezo2* we could measure robust poking-induced currents, the amplitudes of which increased with increasing indentation strength (Figure 1A,B). Similarly, we could record robust inward currents in outside out excised patches to fast pressure pulses taken from N2A^*Piezo1*-/-^ cells expressing *Piezo1*, but could measure no currents in patches taken from cells transfected with *tmem150c* expression constructs or vector transfected control cells (Figure 1C,D). Finally, we plated N2A^*Piezo1*-/-^ cells on elastomeric pillar arrays and evaluated whether pili movement could evoke mechanosensitive currents in cells expressing *tmem150c*. As positive control we used cells expressing *Trpv4 or Piezo2* in which we could measure robust inward currents to pili deflections in a range from 5 - 1000 nm. In contrast, we could measure no deflection-induced current from cells expressing *tmem150c* (Figure 1E,F). We conclude that TMEM150c by itself is unlikely to form a membrane channel that can be gated by force applied through membrane stretch or via extracellular matrix.

**Figure 1.**
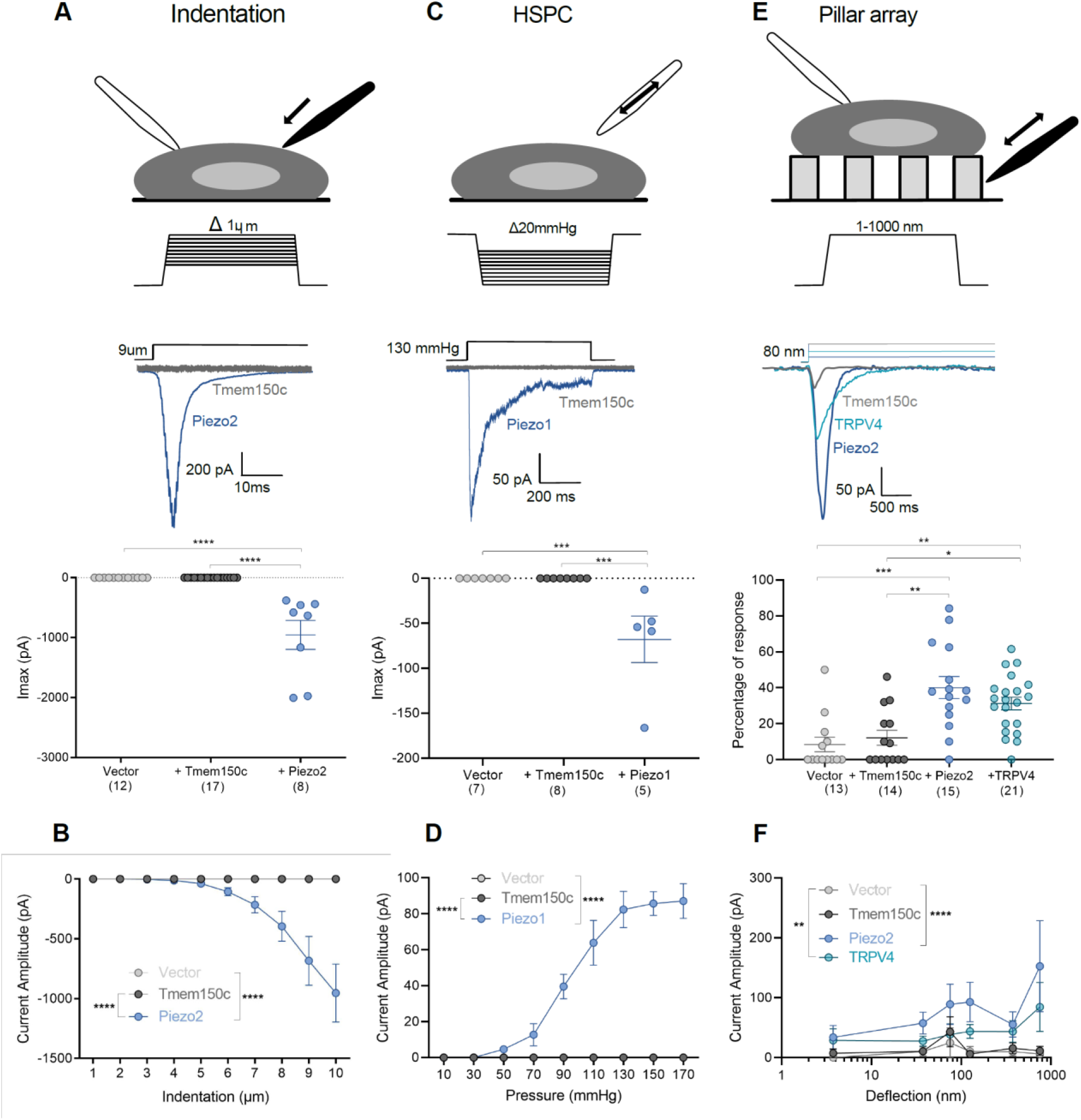
Overexpression of *Tmem150c* in N2A^Piezo1-/-^ cells does not evoke mechanosensitive currents. **(A-C,E)**. Top, cartoons of the different mechanical stimuli applied in this study together with representative traces from mechanosensitive currents evoked using indentation (A), HSPC (C) or pillar arrays (E) methods. Piezo and TRPV4 channels were used as positive controls. Cells overexpressing *Tmem150c* did not display mechanosensitive currents in any of the assays that differed from cells transfected with the empty expression vector. **(A,C,E),** quantified for all stimulus strengths in B,D,F. (A,B) Scatter plots showing the maximal peak (Imax) from mechanosensitive currents observed in N2a P1KO cells overexpressing a vector, Tmem150c or Piezo2 channels. No mechanosensitive currents were observed for N2A^Piezo1-/-^ overexpressing *Tmem150c* (Kruskal-Wallis test **** p<0.0001) with indentation (A,B) or HSPC (C,D) Kruskal-Wallis test **** p<0.0001. N2A^Piezo1-/-^ cells transfected with vector controls occasionally displayed a pillar induced mechanosensitive current, but the frequency of such currents was not higher in cells expressing *Tmem150c* (E). In contrast, almost cells overexpressing either *Piezo2* or *Trpv4* showed large and robust mechanosensitive currents to pillar deflection (E,F) Current kinetics observed in N2A^Piezo1-/-^ overexpressing a vector, TMEM150c or PIEZO2 channels. Left, indentation-current amplitude relationship (A) showing that TMEM150c is indentation-insensitive. One way ANOVA**** p<0.0001. (C) Overexpression of *Tmem150c* was not associated with the pressure activated currents in membrane patches subjected to pressure pulses (C,D) Kruskal-Wallis test **** p<0.0001. Current amplitude-deflection relationship (I) for currents recorded in cells expressing *Tmem150c, Piezo2 or Trpv4*. (F) Cells expressing Tmem150c exhibited similar amplitude-deflection currents as cells transfected with the empty vector Two way ANOVA **** p<0.0001, ** p=0.002.

Hong and colleagues used a mouse model in which the *Tmem150c/Ttn3* gene had been ablated using the Knockout-first strategy (Tate and Skarnes, 2011). They recorded fewer mechanosensitive currents with slowly inactivating properties in *Tmem150c* mutant sensory neurons compared to controls (Hong et al., 2016). We obtained the same mouse mutant strain, but derived from a different ES cell clone, to evaluate the function of TMEM150c in somatosensory neurons. In these mice a β-galactosidase cassette (LacZ) with a splice acceptor was inserted between exon 5 and 6 which should lead to the production of an aberrant truncated transcript encoding a LacZ reporter (Figure 2A). We thus named this allele *Tmem150c^lacZ^* (named *Ttn3^LacZ^* in Hong et al., 2016) and subsequently generated homozygous *Tmem150c^LacZ/LacZ^* on a C57Bl/6N background together with wild type littermate controls (Figure 2). There was no indication that *Tmem150c^LacZ/LacZ^* mice showed embryonic or post-natal lethality and homozygous mice appeared normal and healthy. Since the TMEM150c protein is expressed in almost all sensory neurons of the DRG (Hong et al., 2016) we chose to first evaluate whether loss of TMEM150c is associated with a quantitative change in the receptor properties of cutaneous sensory fibers, which make up the majority of the DRG. We used an ex vivo skin nerve preparation to record from single identified mechanoreceptors and nociceptors in the saphenous nerve using recording and stimulation protocols previously described (Schwaller et al., 2021; Walcher et al., 2018; Milenkovic et al., 2008). In total we made recordings from over 100 single fibers in wild type and *Tmem150c^LacZ/LacZ^* mice (Table 1). The proportions of A-β-fibers that could be classified as rapidly-adapting mechanoreceptors (RAMs) or slowly-adapting mechanoreceptors (SAMs) was unchanged in *Tmem150c^lacZ/LacZ^* mice compared to controls. Evaluation of the stimulus response properties of both mechanoreceptors and nociceptors revealed no significant differences between the two genotypes (data not shown). These negative results led us to evaluate whether there had been efficient ablation of the *Tmem150c* gene in this mutant mouse. TMEM150c is widely expressed, but public gene expression databases indicated that its is especially strongly expressed in epididymal tissue (Yue et al., 2014). Under ideal circumstances staining for β-galactosidase activity should reveal the cells that endogenously express the *Tmem150c* transcript in *Tmem150c^LacZ/LacZ^* mice. Indeed, we could observe robust β-galactosidase expression in the cells of the epididymis, however, we were never able to detect a signal in sections taken from the DRG (Figure 2B). This prompted us to ask whether there was splicing around the β-galactosidase neomycin cassette in sensory neurons. Using RT-PCR for an amplicon located in the *Tmem150c* transcript downstream of the intronic neomycin insertion we could indeed amplify a PCR product in the DRG, but not in the epididymis of *Tmem150c^LacZ/LacZ^* mice (Figure 2C). We cloned and sequenced the amplicon amplified from the DRGs of *Tmem150c^LacZ/LacZ^* mice and could confirm that the sequence was identical to the mature *Tmem150c* transcript. Thus, at least some full-length transcripts encoding the TMEM150c protein is present in the DRGs of *Tmem150c^LacZ/LacZ^* mice. These data indicated that the genetic strategy for ablating the *Tmem150c* gene using the knockout-first allele may not lead to complete ablation of TMEM150c at least in sensory neurons.

**Figure 2.**
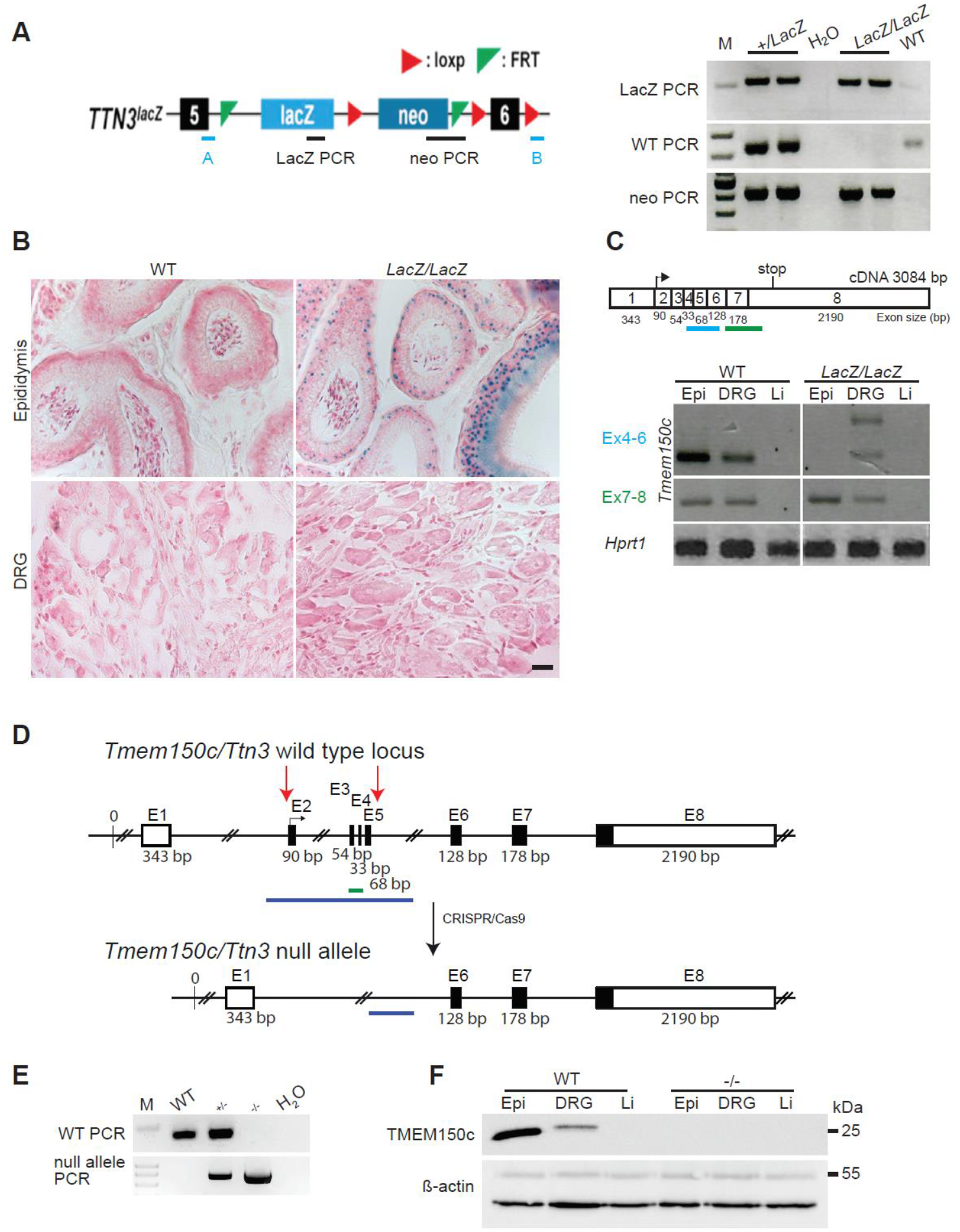
A) Left panel: Schematic representation of *Tmem150c/Ttn3* locus of the knockout first mouse (*Ttn3^LacZ^*) named here *Tmem150c^LacZ^* generated by the trans-National Institutes of Health Mouse initiative knockout Mouse project (https://www.komp.org) containing a LacZ cassette and a neomycin cassette with a stop codon inserted between exons 5 and 6 and resulting in a frame shift. Right panel: Representative gels showing PCR results for neomycin cassette, LacZ cassette and wild type (WT) bands. A and B primers (in blue) used for the amplification of the WT band. Genomic DNA from ear biopsies from WT, heterozygous and homozygous mice were analysed. B) β-galactosidase staining (blue color) of epididymis (positive tissue control) and DRG of *Tmem150c^LacZ/LacZ^* mice. Note the unexpected lack of staining in the DRGs of *Tmem150c^Lacz/LacZ^* mice. WT tissues used as negative control. Scale bar = 20 μm. C) Top panel: schematic representation of *Tmem150c* cDNA (source ensembl Tmem150c-201) containing 8 exons (1-8) with the start codon located in exon 2 (2) and the stop codon located in exon 8 (8). Bottom panel: RT-PCR performed on cDNA prepared from tissues of WT mice and *Tmem150c^Lacz/LacZ^* mice with the blue line indicating amplicon covering exon 4 to 6 (the targeted area of *Tmem150c^LacZ^* allele) and the green line indicating amplicon covering end of exon 6 to beginning of exon 8. Note the presence of unexpected bands in DRG of the *Tmem150c^Lacz/LacZ^* mice but the absence of bands in Epi and Li as expected. *Hprt1* (housekeeping gene) is used as positive control for all tissues. Epi: epididymis (positive control), DRG: dorsal root ganglia, and Li: liver (negative control). D). Schematic representation of *Tmem150c/Ttn3* knock-out (-/-) locus using CRISPR/Cas9 technology: deletion of nucleotide sequence between end of intron 1-2 and beginning of intron 5-6. WT allele and null allele (KO) are shown: Exon1 (E1) encodes the 5’UTR and E8 the 3’UTR (white box). Black box: coding sequence. (source ensembl ENSMUSG0000005064017). Red arrows indicate the location of gRNA sequences used for CRISPR/Cas9. E) PCR performed on genomic DNA from ear biopsies from WT, heterozygous and homozygous mice are shown. The WT amplicon is represented by the green line in the scheme covering E3 and E4 (179 bp). The null allele amplicon is represented by the blue line producing a 886 bp fragment when the nucleotide sequence between intron 1-2 and intron 5-6 (3039 bp in WT) is deleted as in the null allele. Note that this PCR fragment could amplify a 3039 bp fragment in WT mice (see blue line) but this is not possible using the chosen PCR conditions. M: DNA marker. F) TMEM150c protein expression using Western blot showing absence of TMEM150c protein in tissues from *Tmem150c*^-/-^ mice. β-actin was used as loading control.

**Table 1.**
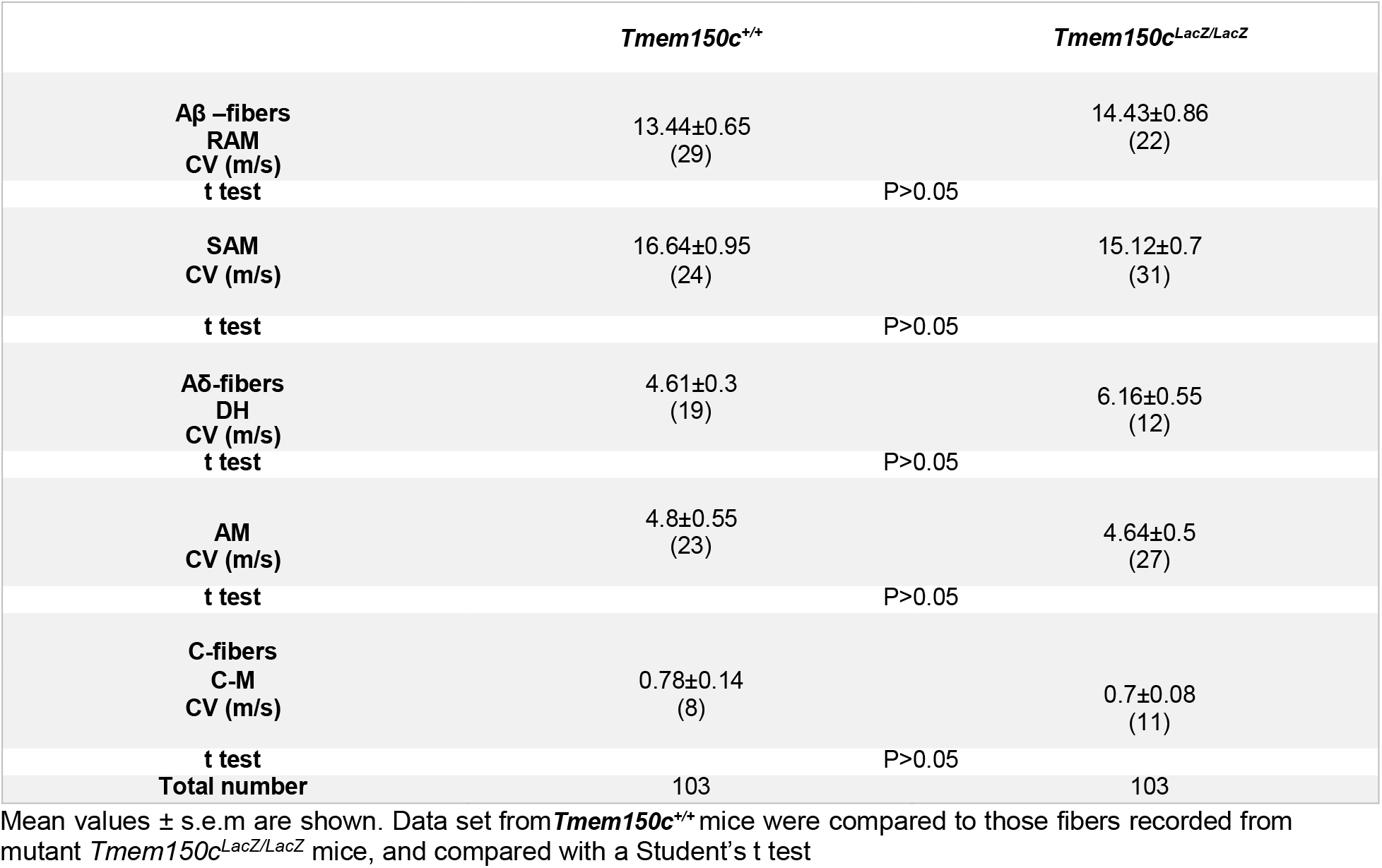
Proportions and conduction velocities of sensory afferents recorded from *Tmem150c^LacZ/LacZ^* from saphenous skin-nerve preparation.

In order to reliably examine the functional importance of TMEM150c in sensory neurons we generated another allele in which the *Tmem150c* locus was ablated using a CRISPR/Cas9 gene editing strategy (Figure 2). In this strategy a genomic region encompassing Exon2-Exon5 was excised so that the sequences encoding transmembrane domains 1-2 were deleted followed by a frameshift. We obtained live heterozygous mice with this allele which we believe is a true null allele (*Tmem150c*^-^). We were able to generate *Tmem150c*^-/-^ mice which were also appeared viable and healthy, like *Tmem150c^LacZ/lacZ^* mice. We could confirm expression of the TMEM150c protein in wild type mouse DRGs using Western blotting which revealed a specific protein band with an apparent molecular weight of ca 25 kDa as predicted from the amino acid sequence. Importantly the TMEM150c protein band was completely absent from the DRG of *Tmem150c*^-/-^ mice, confirming that these mice are true null mutants (Figure 2). It is possible that loss of TMEM150c may affect the development or anatomical integrity of the peripheral nervous system. We thus made an anatomical assessment of the peripheral nerves of wild type and *Tmem150c*^-/-^ mutant mice. Using transmission electron microscopy we examined the ultrastructure of sensory axons in the tibial nerve and found normal large and thinly myelinated axons. We calculated the G-ratios for the myelin sheath which was not different between wild type *Tmem150c*^-/-^ mice. Unmyelinated fibers were present in Remak bundles also had normal morphologies in *Tmem150c*^-/-^ mice. Finally, we counted the numbers of myelinated and unmyelinated axons in the peripheral nerve and found no evidence of axonal loss in *Tmem150c*^-/-^ mice (Figure 3).

**Figure 3.**
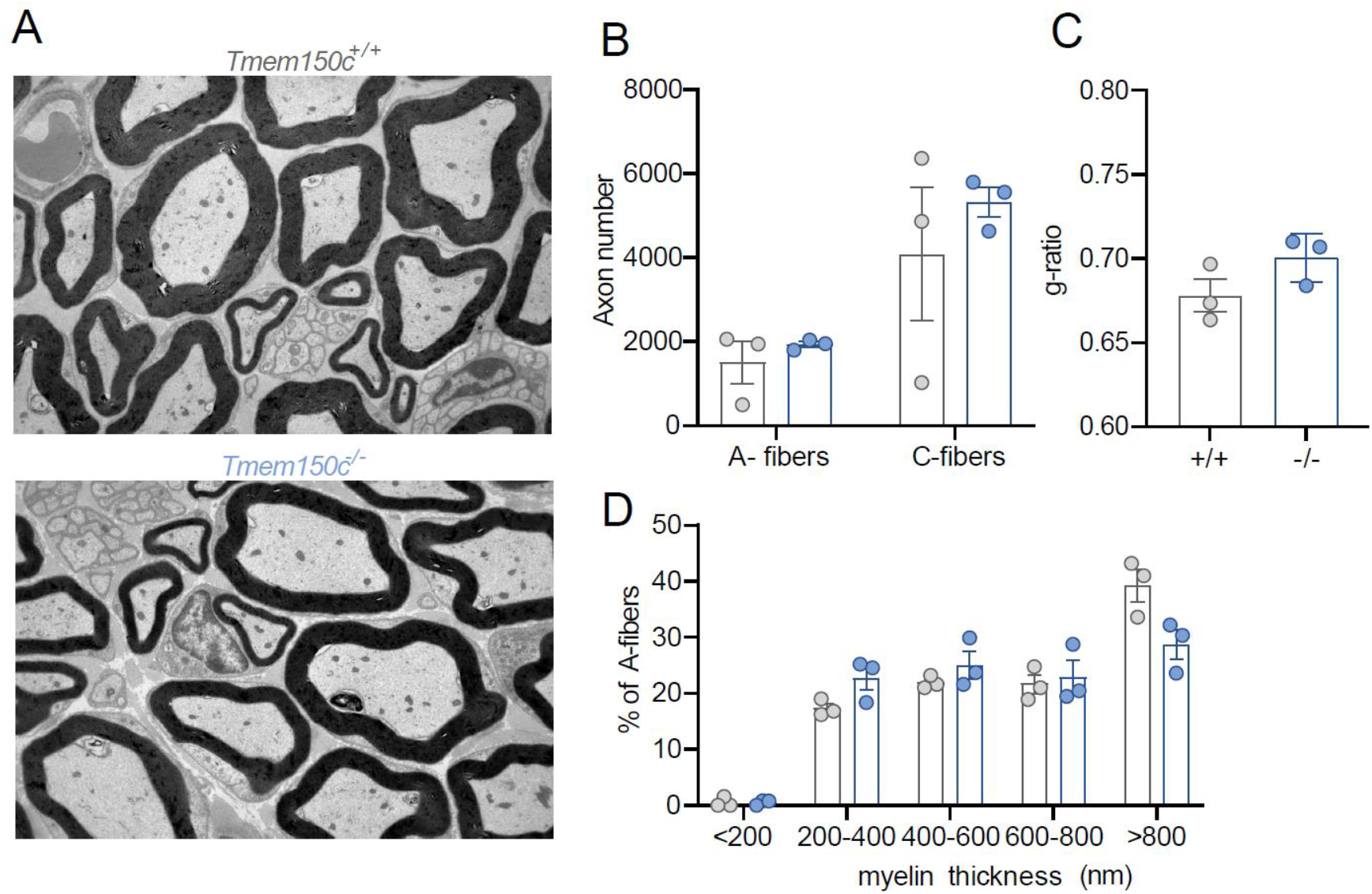
Ultrastructural analysis of the Tibial nerve. A. Example transmission electronmicrographs from the tibial nerve of a wild type *Tmem150c^+/+^* (top) and *Tmem150c*^-/-^ mouse (bottom). B. Quantification of axon numbers revealed no loss of myelinated A-fibers or unmyelinated C-fibers. Also measurements of the G-ratio a measure of myelin thickness showed no difference between genotypes. C. Quantification of myelin thickness across all axon sizes (binned by mylin thickness) revealed no differences between nerves taken from control *Tmem150c*^+/+^ and *Tmem150c*^-/-^ mice.

Although TMEM150c seems unlikely to be able to form a mechanosensitive ion channel by itself (Figure 1), it may have an important modulatory function on components of the mechanotransduction apparatus in sensory neurons. We thus examined in detail the physiological properties of mechanoreceptors and nociceptors in the tibial nerve that innervate the glabrous skin of the mouse hindlimb in wild type and *Tmem150c*^-/-^ mice using established methodology (Walcher et al., 2018; Schwaller et al., 2021). We recorded from at least 40 single Aβ-fiber mechanoreceptors in each genotype and noted no change in the proportion of fibers that could be classified as RAMs or SAMs (Table 2). Two types of mechanoreceptors, Aβ-fiber RAMs and Aδ-fiber D-hair receptors, only respond to moving mechanical stimuli and show distinctive velocity sensitivity that can be measured using a series of ramp stimuli with increasing velocity (Heidenreich et al., 2012; Walcher et al., 2018; Schwaller et al., 2021). RAMs and D-hair receptors in *Tmem150c*^-/-^ had typically low mechanical thresholds that did not differ from those in wild type mice (Figure 4A,B). The velocity sensitivity of these two mechanoreceptors were also indistinguishable between wild type and *Tmem150c*^-/-^ mice (Figure 4C,D).

**Table 2.** Proportions and conduction velocities of primary afferents recorded from *Tmem150*^-/-^ compared to *Tmem150c^+/+^* mice.

|  | *Tmem150c*<sup>+/+</sup> | *Tmem150c*<sup>-/-</sup> |
| --- | --- | --- |
| **Aβ –fibers**<br>**RAM**<br>**CV (m/s)** | 14.28±1.8<br>(20) | 12.29±1.53<br>(24) |
| **t test** | P>0.05 |  |
| **SAM**<br>**CV (m/s)** | 12.41±1.43<br>(26) | 11.81±1.98<br>(32) |
| **t test** | P>0.05 |  |
| **Aδ-fibers**<br>**DH**<br>**CV (m/s)** | 6.93±1.03<br>(16) | 6.66±1.47<br>(18) |
| **t test** | P>0.05 |  |
| **AM**<br>**CV (m/s)** | 5.35±1.47<br>(16) | 3.89±1.42<br>(25) |
| **t test** | P>0.05 |  |
| **C-fibers**<br>**CV (m/s)** | 0.56±0.07<br>(26) | 0.68±0.11<br>(24) |
| **t test** | P>0.05 |  |
| **Total number** | 104 | 123 |

Student's T-test was used to test for differences between recordings made from wild type and *Tmem150*<sup>-/-</sup> mice

**Figure 4.**
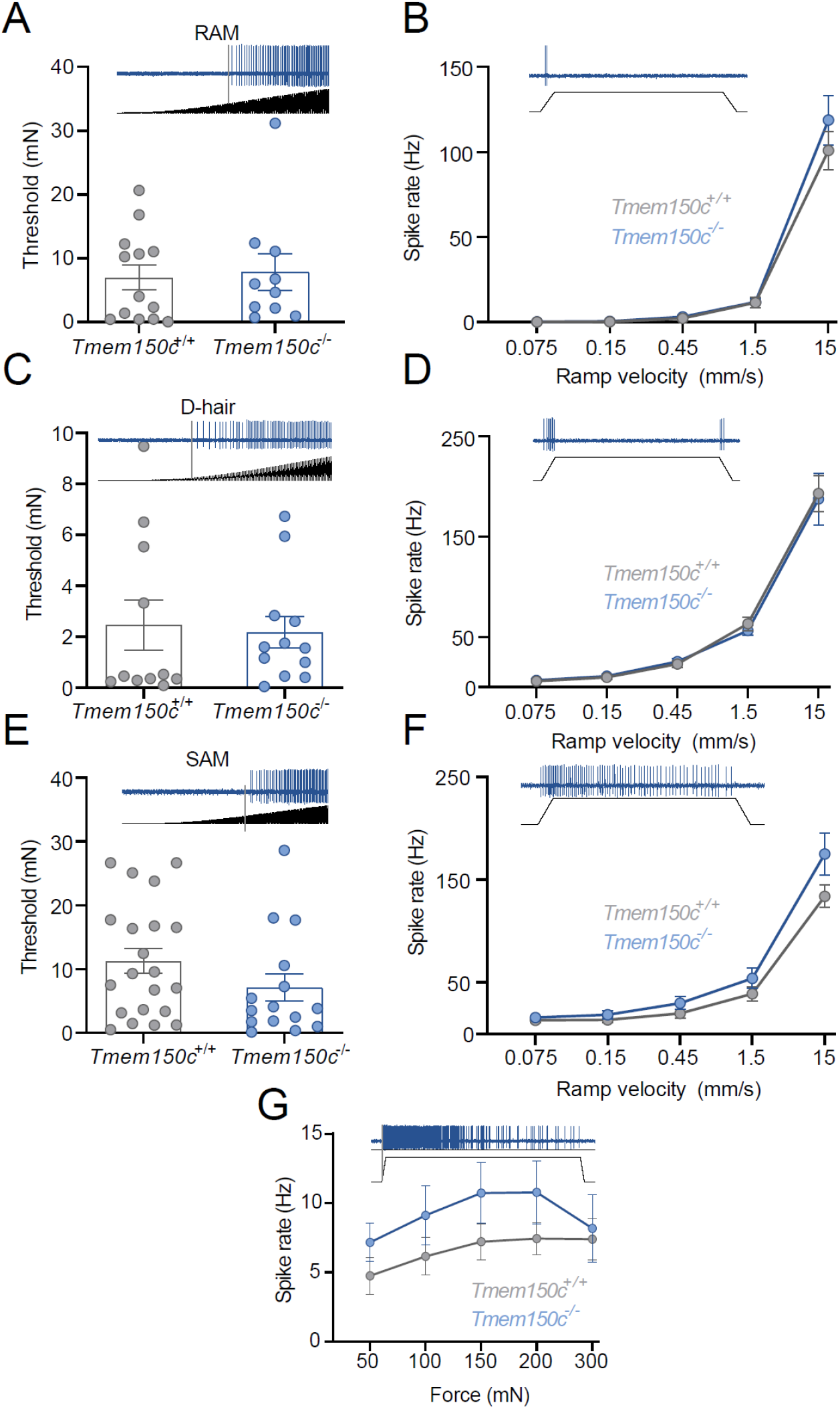
Low threshold mechanoreceptors are unchanged in *Tmem150*^-/-^ mice. Rapidly-adapting mechanoreceptors were stimulated with a linearly increasing 5 Hz sinusoidal force stimulus to determine mechanical threshold in mN (A) or a series of ramp and hold stimuli of different velocities (B) (example recordings shown). There was no difference in these mean parameters in RAMs recorded from *Tmem150*^-/-^ mice compared to littermate controls (*Tmem150c*^+/+^ mice). D-hair mechanoreceptors were also probed with the same quantitative stimuli (C, D) There was no difference in the mean mechanical threshold or velocity sensitivity of D-hair mechanoreceptors between the two genotypes. Slowly-adapting mechanoreceptors (SAMs) were also probed with the same quantitative stimuli (E,F), as well as a series of 10s long ramp and hold stimuli of increasing holding force (50-300 mN) (G). There was no statistical difference in the mean mechanical threshold, velocity sensitivity or stimulus response function of SAMs between the two genotypes. There was a tendency for the mean spike rate to be slightly higher in SAMs from *Tmem150*^-/-^compared to controls (G), but this difference was not statistically significant. The numbers of fibers included in the analysis can be found in Table 2.

In their initial studies Hong and colleagues claimed that TMEM150c could mediate a mechanosensitive current with slow inactivation. Almost all mechanoreceptors display so called RA-mechanosensitive currents that inactivate in less than 5 ms (Hu and Lewin, 2006; Lechner and Lewin, 2013; Lechner et al., 2009). It is thus possible that lack of TMEM150c has no effect of mechanoreceptors that normally only display RA-mechanosensitive currents. We thus studied mechanoreceptors and nociceptors which may possess mechanosensitive currents that inactivate relatively slowly (Hu and Lewin, 2006; Lechner et al., 2009; Lechner and Lewin, 2009). We recorded from both myelinated and unmyelinated cutaneous nociceptors in control and *Tmem150c*^-/-^ mice (Table 2). We used a series of ramp and hold mechanical stimuli to probe their stimulus response properties. Slowly-adapting mechanoreceptors (SAMs) in the glabrous skin innervate Merkel cell complexes and are both velocity sensitive and respond tonically to small static forces. We also found that the velocity sensitivity and static force sensitivity of SAMs in *Tmem150c*^-/-^ mice were statistically indistinguishable from those recorded in wild type mice (Figure 4). The firing of SAMs to suprathreshold stimuli in *Tmem150c*^-/-^ mice was on average slightly elevated compared to those in control animals, but this apparent difference was not statistically different (Figure 4).

We measured force required to evoke the first spike during the ramp of the stimulus (Figure 5A) and found that for AMs the mean threshold for activation in *Tmem150c*^-/-^ mutant mice was almost identical to those measured from wild type (Figure 5B). The stimulus response function of AMs in wild type and *Tmem150c*^-/-^ mice were also not different (Figure 5C). We divided C-fiber nociceptors into those that only respond to mechanical stimuli and do not respond to heating and cooling stimuli, so called C-mechanonociceptors (CMs) and Polymodal nociceptors, which respond to mechanical and thermal stimuli. These two types of nociceptors have different stimulus response properties to mechanical force (examples in Figure 5A) (Milenkovic et al., 2008). Neither C-Ms nor Polymodal nociceptors showed different mean force thresholds or stimulus response properties in *Tmem150c*^-/-^ mice compared to controls (Figure 5 D-G). We conclude that the mechanosensory properties of virtually all types of cutaneous sensory neurons is unaffected in the absence of TMEM150c.

**Figure 5.**
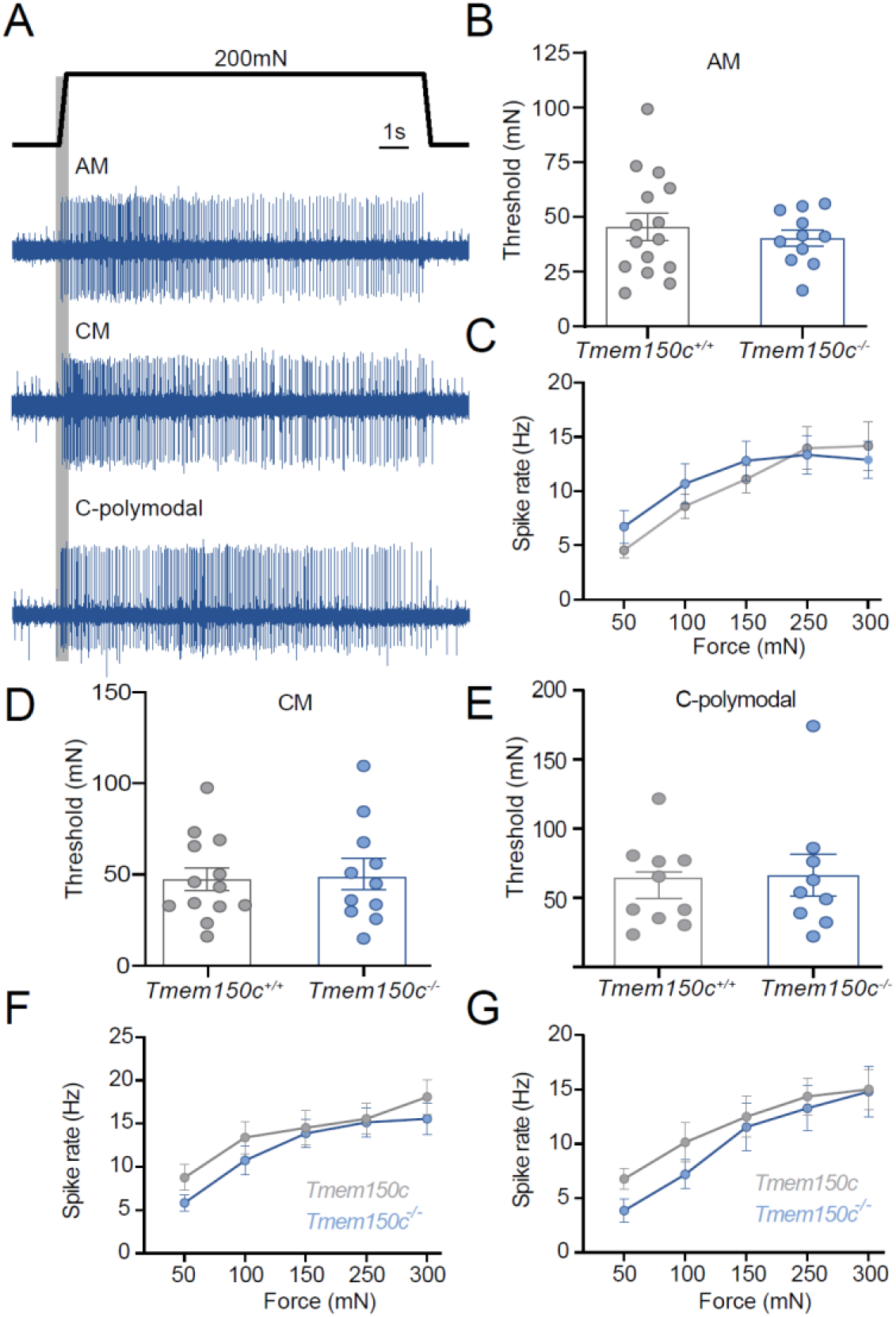
The mechanosensitivity of nociceptors was unchanged in *Tmem150*^-/-^ mice. We recorded from thinly myelinated mechanonociceptors termed A-mechanonociceptors (AM), C-fiber mechanonociceptors (CM) and polymodal C-fiber nociceptors, the latter respond to mechanical stimuli and at least one other stimulus modality like heat or cold. A, Example recordings from these three receptor types in response to a ramp and hold controlled force stimuli. Mechanical thresholds were measured as the force required to evoke the first spike during the stimulus ramp phase. B, For AMs, there was no difference in the mean mechanical threshold between fibers recorded from *Tmem150*^-/-^ mice compared to littermate controls (*Tmem150c*^+/+^ mice). Stimulus response properties of AMs to controlled force stimuli were also not different between genotypes (C). E, F, The mean mechanical thresholds of CMs and C-polymodal fibers did not differ between genotypes. The stimulus response functions of CM and C-polymodal fibers also did not differ between genotypes F,G. The numbers of fibers included in the analysis can be found in Table 2.

In their original paper examining mice with a *Tmem150c* allele generated through the knockout first strategy Hong and colleagues described a deficit in motor coordination in these mice that they attributed to changes in proprioceptor properties (Hong et al., 2016). We did not notice any overt motor coordination deficits in *Tmem150c*^-/-^ mice, but decided to quantify motor behavior using a so-called catwalk assay (Mendes et al., 2015). We analyzed various gait parameters such as locomotion speed, regularity of swing phase and step distances and could not detect any significant deficits in *Tmem150c*^-/-^ mice compared to wild type controls (Figure 6).

**Figure 6.**
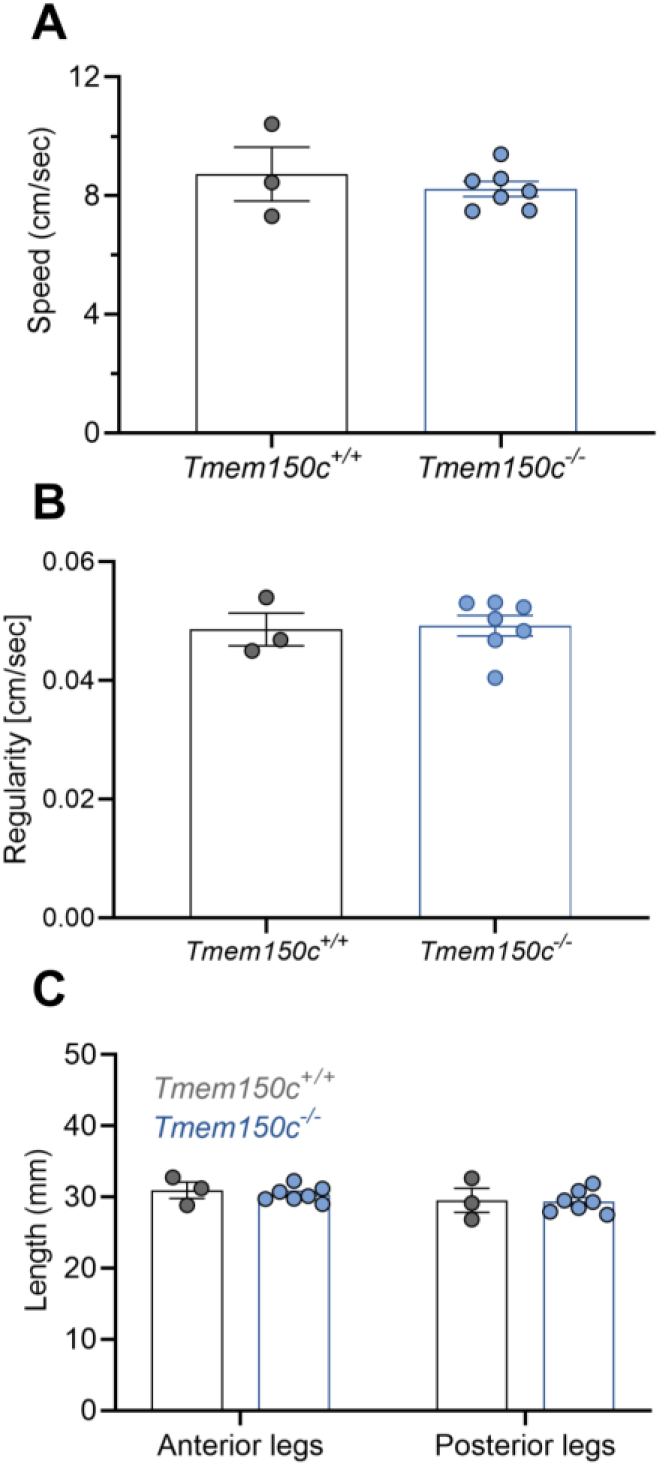
Mouse walk behavior test shows comparable motor coordination in *Tmem150c*^-/-^ and *Tmem150c*^-+/+^ mice. (A) Average speed of each mouse from one extreme to the other in the walkway, showing no differences between *Tmem150c*-+/+ mice (8.72±0.90) and *Tmem150c*-/- (8.22±0.25) mice. (B) Regularity of swing time for all legs is similar in *Tmem150c*-+/+ mice (0.04±0.002) and *Tmem150c*-/- (0.04±0.001) mice. (B) No differences were observed in step distance between each swings in control *Tmem150c*-+/+ and *Tmem150c*-/- mice; anterior legs (WT 28.86±1.14; KO 29.02±0.41) and posterior legs (WT 26.85±1.6; KO 27.52±0.59). Data expressed as mean ± s.e.m.

## Discussion

The membrane protein TMEM150c or Tentonin3 has been proposed to form a mechanosensitive ion channel in proprioceptors, nodose neurons innervating the aortic arch (Lu et al., 2020) and even in pancreatic β cells (Wee et al., 2021). However, early follow up studies could not reproduce the finding that heterologous expression of *Tmem150c* is associated with the appearance of mechanosensitive currents, when PIEZO1 channels are absent (Anderson et al., 2018; Dubin et al., 2017). Here we could confirm that the expression of *Tmem150c* in N2a cells lacking PIEZO1 channels is not associated with the appearance of indentation evoked currents. In addition, we could show that two further modes of stimulation that efficiently activate mechanosensitive channels, membrane stretch and substrate deflection also do not evoke currents in cells expressing *Tmem150c* (Figure 1). Using a knockout mouse model it has been claimed that loss of the TMEM150c protein is associated with a loss of mechanosensitive currents with slow inactivation kinetics in DRG neurons, nodose neurons and pancreatic β-cells (Hong et al., 2016; Lu et al., 2020; Wee et al., 2021). Here we used the same mouse model, but found that the knockout-first gene targeting strategy was associated with remaining mature wild type *Tmem150c* transcripts in the DRG (Figure 2). We could also not detect β-galactosidase activity in the DRG which should be present in cells in which the gene has been effectively silenced. We concluded that the *Tmem150c^lacZ/lacZ^* mice may not produce a complete gene ablation at least in some tissues. In order to better test for a possible function of TMEM150c in vivo we created a new mouse model in which a large part of the *Tmem150c* locus was ablated. This new *Tmem150c*^-/-^ mouse was viable and seemingly healthy. Even if TMEM150c forms an accessory subunit of mechanosensitive ion channels then it would seem likely that sensory neurons in *Tmem150c*^-/-^ mice would show physiological deficits. We made extensive single-unit recordings from cutaneous sensory neurons from wild type and *Tmem150c*^-/-^ mutant mice. We detected no deficits in the mechanosensitivity of sensory neurons lacking TMEM150c. If *Tmem150c*^-/-^ mutant mice have strongly reduced proprioceptor function we might expect to observe deficits in locomotor behviour. However, an analysis of the gait of *Tmem150c*^-/-^ mutant mice revealed that these animals have normal gait parameters indistinguishable from wild type mice. We conclude that there is little or no evidence that the TMEM150c protein participates in sensory mechanotransduction.

Most of the evidence gathered for a functional role of TMEM150c in mediating endogenous slowly inactivating mechanosensitive currents relies on the use of the *Tmem150c^lacZ/LacZ^* mouse model (Lu et al., 2020; Hong et al., 2016; Wee et al., 2021). Here we show that *Tmem150c^lacZ/LacZ^* mice may carry a hypomorphic mutation that allows expression of wild type TMEM150c in some tissues. The wide range of phenotypes described in this mouse model should therefore be treated with some caution. It is known that insertion of genetic elements into introns which is a necessary part of the knockout first strategy (Tate and Skarnes, 2011) may lead to unexpected effects even on the expression of unrelated genes (Pan et al., 2016). In the published studies using the *Tmem150c^lacZ/LacZ^* mouse model there was no examination of LacZ activity in the cell types studied with reduced mechanosensitive current. Furthermore, proof of the success of the gene deletion appeared to be solely based on lack of immunopositive cells in *Tmem150c^lacZ/LacZ^* tissues (Wee et al., 2021; Lu et al., 2020; Hong et al., 2016). We used antibodies directed against the same epitope described in (Hong et al., 2016) to detect the protein in wild type DRG sections and in sections from *Tmem150c*^-/-^ mice. Unfortunately, in our hands we always observed a substantial background staining in DRG sections taken from *Tmem150c*^-/-^ mice using such antibodies (data not shown).

Here we used the *Tmem150c*^-/-^ mutant mice to try and definitively answer the question of whether TMEM150c has a non-redundant function in sensory neurons of the DRG. It is clear that TMEM150c is found in many DRG neurons (Hong et al., 2016; Parpaite et al., 2021; Usoskin et al., 2015), but recent experiments have also cast doubt on whether the presence of TMEM150c is required for the expression of mechanosensitive currents in cultured sensory neurons (Parpaite et al., 2021). Additionally, there is now evidence that sensory neuron mechanosensitivity may not be solely dependent on channels within the sensory cell membrane. Thus, recently identified specialized Schwann cells that surround the peripheral endings of unmyelinated nociceptors may directly participate in the detection of noxious mechanical stimuli (Abdo et al., 2019). In addition, cells that form specialized sensory end-organs innervated by mechanoreceptors in birds were also recently shown to be mechanosensitive (Nikolaev et al., 2020). Using our global *Tmem150c*^-/-^ mutant mice we wished to test whether loss of the TMEM150c protein had an impact on the mechanosensitive properties on functionally identified sensory neurons using *ex vivo* skin nerve preparations. Recordings from identified sensory neuron populations has in the past proved a reliable way to reveal physiological deficits in mice lacking genes that participate in sensory mechanotransduction or regulate excitability (Heidenreich et al., 2012; Wang and Lewin, 2011; Ranade et al., 2014; Wetzel et al., 2007; Ventéo et al., 2016; Schwaller et al., 2021; Dawes et al., 2018; Murthy et al., 2018). We reasoned that we should observe some deficits in defined sensory neuron types if TMEM150c is required for the expression of slowly inactivating currents in sensory neurons or in sensory Schwann cells. Instead, here we detected no change in the physiological properties of identified nociceptors and mechanoreceptors in mice lacking TMEM150c. It is of course conceivable that we did not observe deficits because the effect was subtle or found in a rare sub-type of mechanoreceptor that was not adequately sampled. In the experiments described here we only characterized the receptor properties of neurons which were first identified with mechanical search stimulus. Sensory fibers lacking a mechanosensitive receptive field, like those found in *Stoml3*^-/-^ or *Piezo2*^-/-^ mutant mice, could have been missed in this study. However, in mutant mice with such a profound mechanosensory deficits we always found deficits in the stimulus response properties of remaining afferent populations (Wetzel et al., 2007; Ranade et al., 2014; Murthy et al., 2018). In their original paper Hong and colleagues claimed to observe deficits in the muscle stretch-activated activity of muscle afferents innervating the extensor digitorum longus muscle in the hind limb (Hong et al., 2016). In their study Hong and colleagues recorded afferent multi-unit activity from the whole nerve (Hong et al., 2016), but did not identify and classify functional sub-types of muscle afferents (Lewin and McMahon, 1991). Here we did not attempt to make a detailed analysis of identified muscle afferents as we did not observe any deficits in gait parameters (Figure 6).

In summary, we have provided comprehensive evidence that TMEM150c does not by itself form a mechanosensitive ion channel. Furthermore, we show that in animals with a true homozygous null mutation of the *Tmem150c* locus there is no indication of any physiological deficits in any of the major physiologically defined mechanoreceptor and nociceptor populations that innervate the skin. We conclude that it is unlikely that the TMEM150c protein is a necessary regulator of mechanosensitive ion channels that are required for the normal function of somatosensory neurons.

## Acknowledgments

This research was funded by the ERC and the Deutsche Forschungsgemeinschaft (Sensational Tethers 789128 to GRL; SFB 958 Project A9). Technical assistance was provided by Maria Braunschweig, Kathleen Barda, Franziska Bartelt and Christina Schiel. We thank members of the Lewin lab for constructive comments on the manuscript.

## Author contributions

Conceptualization: J.O-A, G.R.L and V.B.; mouse models/experimental design: V.B., G.R.L. and J.O-A.; nerve recording and formal analysis: J.O-A: mouse behavior J.O-A and J.A.G.C; patch clamp electrophysiology: J.O-A and J.A.G.C; electron microscopy B.P. J.O-A., A.F.C.P and V.B.; writing G.R.L. and J. O-A. with input from all authors. Supervision and funding: G.R.L.

## Competing interests

Authors declare no competing interests.

## Data and materials availability

All data is available in the main text or the supplementary materials. All data, code, and materials used in the analysis must be available in some form to any researcher for purposes of reproducing or extending the analysis.

## Notes

### Competing Interest Statement

The authors have declared no competing interest.

